# A Systems approach identifies MERTK as a therapeutic vulnerability in ZFTA-RELA-driven ependymomas

**DOI:** 10.1101/2025.02.06.636949

**Authors:** Marina Chan, Songli Zhu, Zach Russell, Sonali Arora, Aleena Arakaki, Joel M Vaz, Deby Kumasaka, Frank Szulzewsky, Antony Michealraj, Eric C Holland, Taranjit S Gujral

## Abstract

Ependymomas (EPN) are rare central nervous system tumors that account for approximately 10% of intracranial tumors in children and 4% in adults. Despite their clinical and molecular heterogeneity, spanning supratentorial, posterior fossa, and spinal subtypes, treatment remains limited to surgery and radiotherapy, with chemotherapy offering minimal benefit. Here, we performed transcriptomic analysis of 370 human ependymoma samples and identified two distinct molecular subgroups: EPN-E1 and EPN-E2. The EPN-E1 cluster is enriched for supratentorial tumors harboring ZFTA-RELA fusions (ZFTA-RELA^fus^), which occur in over 70% of cases and are associated with poor prognosis. To identify targeted therapies for this aggressive subtype, we validated a ZFTA-RELA^fus^ mouse model that recapitulates the human EPN-E1 transcriptome and used it for target discovery. Through Kinome Regularization, a machine learning-driven polypharmacology approach, we identified MERTK as a critical regulator of tumor cell viability. Genetic depletion or pharmacologic inhibition of Mertk reduced cell growth *ex vivo*, and treatment with a clinical-grade MERTK inhibitor significantly suppressed tumor proliferation *in vivo*. Both human EPN-E1 tumors and ZFTA-RELA^fus^ mouse tumors exhibited elevated expression of MERTK and its ligand GAS6, and MERTK inhibition led to suppression of pro-survival signaling pathways including MEK/ERK and PI3K/AKT. Notably, over 80% of genes upregulated in ZFTA-RELA^fus^ tumors were downregulated following MERTK inhibition, indicating a strong dependency on this pathway for tumor maintenance. These findings define a signaling vulnerability in ZFTA-RELA-driven ependymomas and support the clinical development of MERTK-targeted therapies for patients with the high-risk EPN-E1 subtype.

## Introduction

Ependymomas are rare cancers that account for approximately 10% of intracranial tumors in children and 4% of brain and spinal cord tumors in adults(*1*). They arise throughout the central nervous system and include supratentorial, posterior fossa, and spinal subtypes, each representing clinically and molecularly distinct entities. Treatment remains largely limited to surgery and radiation, with chemotherapy offering limited benefit, primarily in select pediatric cases (*2, 3*). Recent genomic studies have uncovered recurrent chromosomal translocations in supratentorial ependymomas, most notably ZFTA-RELA fusions, which occur in over 70% of cases and are associated with poor outcomes. (*3, 4*). Despite growing insights into the molecular landscape of these tumors, there are currently no therapies that directly target the fusion proteins. This highlights a broader challenge in rare cancers, where limited patient numbers and a lack of tailored models hinder therapeutic discovery. To uncover actionable vulnerabilities and accelerate treatment development for rare subtypes like ZFTA-RELA^fus^ ependymomas, it is essential to (1) comprehensively profile a large number of human tumors to identify robust, subtype-specific molecular signatures, (2) develop physiologically relevant mouse models that accurately reflect human disease, and (3) leverage these models to dissect key signaling pathways and uncover therapeutic targets unique to each molecular subtype.

To address this unmet need, we first performed consensus clustering on RNA-seq data from 370 ependymoma tumor samples across five independent studies to define molecular subtypes that could guide model development and therapeutic discovery. This analysis revealed two distinct subgroups: EPN-E1 and EPN-E2. EPN-E1 was composed predominantly of supratentorial tumors with a high frequency of ZFTA-RELA^fus^, while EPN-E2 was enriched for posterior fossa and myxopapillary tumors. Differential gene expression analysis revealed marked transcriptional differences between the two groups, underscoring their biological distinctiveness. To model the rare and aggressive ZFTA-RELA^fus^ subtype, we leveraged a previously established RCAS/tv-a system to express ZFTA-RELA^fus^ in nestin-positive neural cells in mice. The resulting tumors closely mirrored the human EPN-E1 subtype both histologically and transcriptionally, validating this model as a relevant preclinical platform for investigating disease mechanisms and identifying subtype-specific therapeutic vulnerabilities.

To uncover key signaling pathways driving tumor growth in the ZFTA-RELA^fus^ ependymoma subtype (EPN-E1), we leveraged our human aligned-mouse model system and applied Kinome Regularization (KiR), a machine learning-based, systems polypharmacology approach we previously developed to identify kinases essential for specific phenotypes. Using this method in the ZFTA-RELA-driven mouse model, we identified MERTK as a critical kinase uniquely required for the survival and proliferation of ZFTA-RELA^fus^ ependymoma cells. Genetic knockdown or pharmacological inhibition of Mertk significantly impaired cell growth *in vitro* and reduced viability in tumor slices. Treatment with a clinical-grade MERTK inhibitor also reduced tumor growth and proliferation *in vivo*. Consistent with these findings, both MERTK and its ligand Gas6 were significantly upregulated in ZFTA-RELA^fus^ human tumors compared to other ependymoma subtypes. Mechanistically, ZFTA-RELA^fus^ expression was required for maintaining Mertk and Gas6 expression, and inhibiting this axis suppressed pro-survival pathways, including MEK/ERK and PI3K/AKT. These results suggest that ZFTA-RELA^fus^ reprograms signaling dependencies, creating a vulnerability to MERTK inhibition. Our findings elucidate the molecular drivers of this aggressive ependymoma subtype and support the development of MERTK-targeted therapies for clinical translation.

## Results

### Molecular subgrouping of ependymoma uncovers distinct transcriptomic and gene fusion profiles

To define molecular subtypes of human ependymoma and better understand their biological diversity, we analyzed bulk RNA-seq data from 370 tumor samples collected across five independent studies from North America and Europe (*5–9*) (**Table S1**). These samples included 134 supratentorial, 135 posterior fossa, 77 ependymoma (NOS), 11 anaplastic, 9 myxopapillary, and 4 spinal ependymomas. Raw sequencing reads were aligned to the human genome (hg38), and gene counts were generated for all protein-coding genes. Batch effects were corrected using ComBatSeq (R package *sva*), and data were normalized using variance stabilizing transformation (VST). Consensus clustering of the normalized expression data revealed two distinct molecular subgroups: EPN-E1 and EPN-E2 (**Fig. 1A**). EPN-E1 is predominantly composed of supratentorial ependymomas (ST-EPNs), accounting for 77% (97/126) of the samples, followed by 11% (14/126) ependymoma NOS, 5.55% (7/126) PF-EPNs, 3.96% (5/126) Anaplastic EPNs and 2.38% (3/126) spinal EPNs. The EPN-E2 is primarily dominated by PF-EPNs (52.47%,.127/242) followed by 15.28% (37/242) ST-EPNs, 25% (62/242) ependymoma NOS samples, 3.71% (9/242) myxopapillary EPNs and 2.47% (6/242) Anaplastic EPNs (**Fig. S1**). Notably, all myxopapillary tumors were exclusive to the EPN-E2 cluster, while spinal and anaplastic tumors were distributed across both groups. Although some diagnostic labels may not align with the 2021 WHO classification, we retained the original annotations for consistency and transparency.

**Fig. 1.**
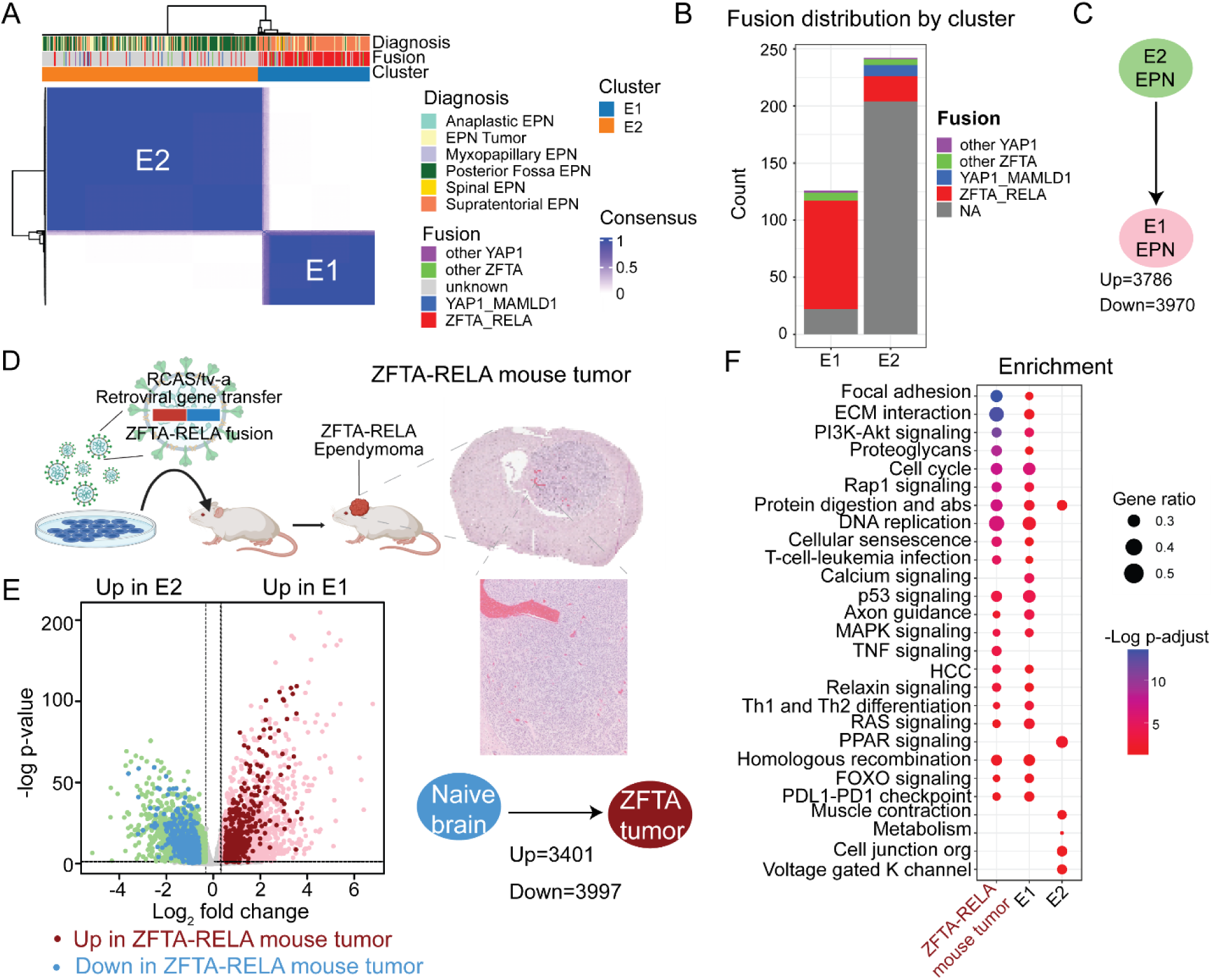
Transcriptomic subtyping of human ependymoma and cross-species alignment with a ZFTA-RELA^fus^ mouse model. **A.** *Human ependymoma subtyping*. Consensus clustering of bulk RNA-seq data from 370 human ependymoma tumors reveals two distinct molecular subgroups, EPN-E1 and EPN-E2.**B.** Barplot showing the distribution of gene fusions across both clusters; EPN-E1 is predominantly composed of ZFTA-RELA^fus^ tumors. **C.** Differential expression analysis between EPN-E1 and EPN-E2 identifies 3,786 upregulated and 3,970 downregulated genes in EPN-E1. **D.** *Development of the ZFTA-RELA^fus^ mouse model.* A schematic depicting the introduction of the ZFTA-RELA^fus^ construct in DAF-1 cells, the resulting tumor development in mice, and histological sections of the mouse brain with ZFTA-RELA^fus^ tumor. Right, schematic showing number of up and down-regulated genes in ZFTA-RELA^fus^ mouse tumor compared to naïve brain. **E.** Volcano plot showing the up-regulated genes (pink) and down-regulated genes (green) in E1 human samples, compared to E2 ependymoma human samples. Up-regulated and downregulated genes in ZFTA-RELA^fus^ mouse tumor compared to naïve brain are shown in red and blue respectively. **E**. Dotplot showing pathways up-regulated in ZFTA-RELA^fus^ mouse tumors compared to Naïve brain, and up-regulated pathways in E1 and E2 human ependymoma samples.

Next, we examined the molecular features associated with each cluster. Analysis of gene fusion profiles revealed that the EPN-E1 group was predominantly composed of supratentorial ependymomas, with 75% of cases harboring ZFTA-RELA^fus^ (**Fig. 1B**). In contrast, the EPN-E2 cluster showed greater molecular heterogeneity and lacked a dominant fusion event. Differential gene expression analysis between the two clusters identified 3,768 genes upregulated and 3,970 genes downregulated in EPN-E1 relative to EPN-E2 (**Fig. 1C**, Supp Table 2a-b), highlighting distinct transcriptional programs associated with each subgroup.

### A ZFTA-RELA-driven mouse model faithfully mirrors the human EPN-E1 subtype

Previous studies, including our own, have demonstrated that the expression of the ZFTA-RELA^fus^ in nestin-expressing brain cells is sufficient to induce EPN in mice(*4, 10, 11*). To model this, we utilized the RCAS/tva system for postnatal cell-type-specific gene transfer(*10*). Specifically, an RCAS retroviral vector encoding the ZFTA-RELA^fus^ was used to infect nestin-expressing cells in *Nestin-tva* mice. This was achieved through intracranial injection of DF1 chicken fibroblasts, transfected to produce the vector. Tumor development occurred within 70-120 days, and affected mice were identified based on clinical signs (e.g., weight loss, poor grooming) and MRI-confirmed intracranial tumors. Tumors were harvested and processed for transcriptomic profiling (**Fig. 1D**). Differential gene expression analysis between ZFTA-RELA^fus^ tumors and naïve brain tissue revealed 3,401 upregulated and 3,997 downregulated genes, confirming that the model recapitulates broad transcriptional reprogramming.

To evaluate its relevance to human disease, we compared the differentially expressed genes from the mouse model to those distinguishing the EPN-E1 and EPN-E2 human subtypes. Genes upregulated in the ZFTA-RELA^fus^ mouse tumors closely aligned with those enriched in the human EPN-E1 cluster, while downregulated genes mirrored those suppressed in EPN-E1 or elevated in EPN-E2 (**Fig. 1E**). Pathway enrichment analysis further showed that biological pathways activated in the mouse model were also specifically enriched in EPN-E1 human tumors, but not in EPN-E2 (**Fig. 1F**). These findings suggests that the ZFTA-RELA^fus^ mouse model is a faithful representation of the EPN-E1 subtype and highlight the central role of ZFTA-RELA in driving tumor development.

### Kinome regularization identifies key kinase dependencies in ZFTA-RELA^fus^ ependymoma

To identify kinases critical for the growth of ZFTA-RELA^fus^ ependymoma cells, we applied our systems polypharmacology approach, Kinome Regularization (KiR), to cancer cells derived from a ZFTA-RELA^fus^ mouse tumor line (referred to as ZR1) (**Fig. 2A**). KiR leverages the polypharmacology of kinase inhibitors, most of which target multiple kinases, to deconvolute kinase dependencies using machine learning. This approach has been successfully applied across diverse biological contexts, including identifying kinases that regulate cell migration(*12–14*), prostate cancer cell growth(*15*), SARS-CoV-2-induced cytokine release in monocytes(*16*), liver-stage malaria(*17*), herpes virus(*18*), and cytokine secretion(*19*), demonstrating its broad utility in uncovering key signaling drivers. Here, we screened a computationally selected training set of 37 kinase inhibitors that together cover over 90% of the human kinome(*13, 15*). Each inhibitor was tested across eight concentrations in ZR1 cells, and drug effects were quantified using a real-time imaging-based growth assay. Staurosporine, a broad-spectrum kinase inhibitor, was included as a positive control (**Fig. S2, Table S2**). The screen generated 73,728 microscopy images, which were analyzed to quantify drug-induced changes in cell viability.

**Fig. 2.**
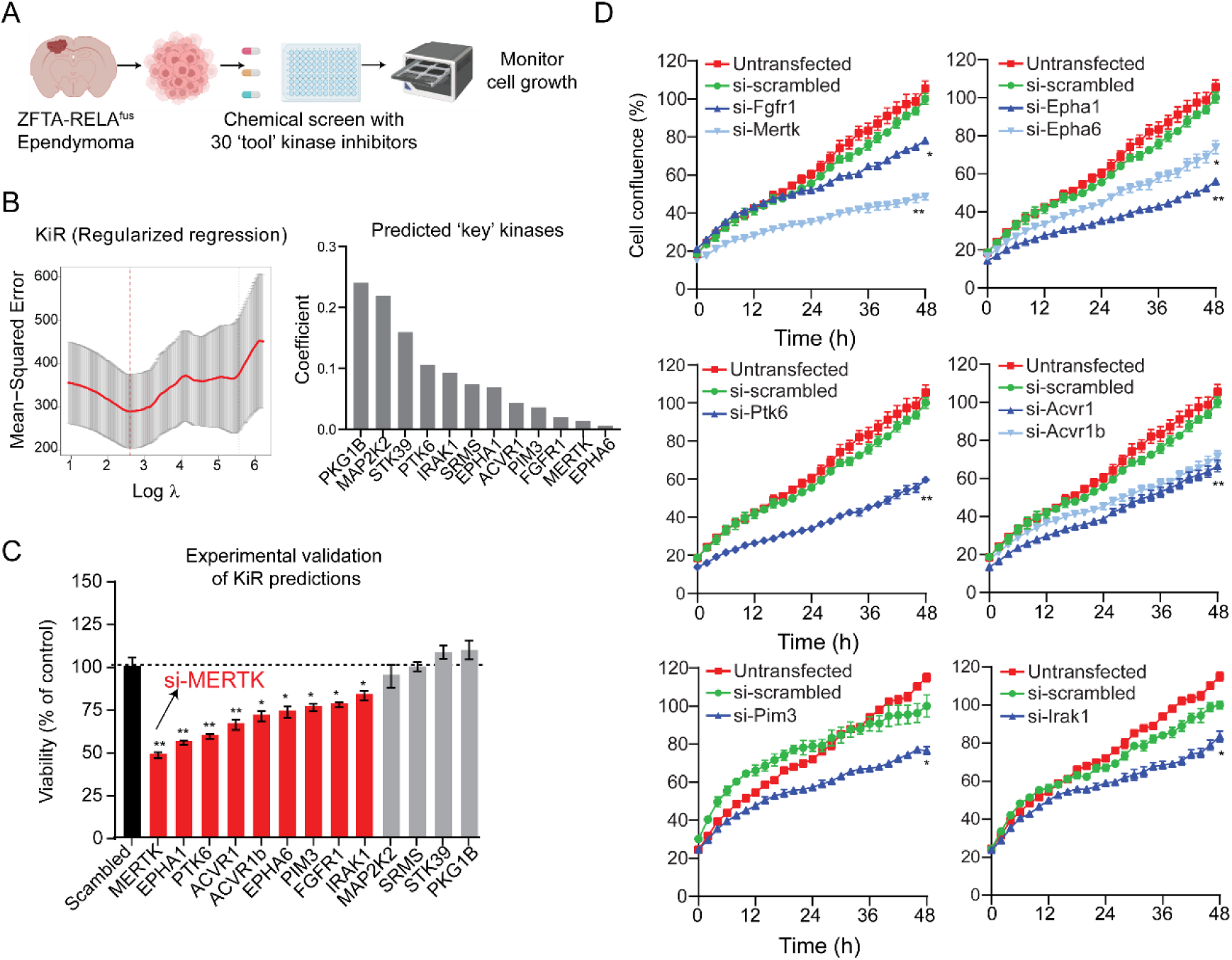
KiR screening identified ‘key’ kinases in ZFTA-RELA^fus^ ependymoma. **A.** A schematic of our experimental strategy for predicting kinases important for the growth of ZFTA-RELA^fus^ - expressing mouse ependymoma cells. A set of computationally chosen kinase inhibitors were evaluated for their effect on the growth of ZFTA-RELA^fus^-expressing cells. Each compound was tested at eight different concentrations. The effect on cell growth was assessed through a real-time growth assay. **B**. KiR regularized regression plot for predicting response to kinase inhibitors in ZR1 cells. The horizontal axis represents the logarithm of the penalty hyperparameter λ, while the vertical axis shows the mean squared error from leave-one-out cross-validation (LOOCV) (mean ± SD, n = 36 folds). The dotted red line indicates the lowest error and the selected value of λ for the final model. **C.** The KiR modeling identified 12 most “informative kinases” out of over 300 kinases that influence the proliferation of ZR1. **D.** Experimental validation of KiR-predicted kinases showed that knockdown of 9 out of 12 kinases resulted in a statistically significant decrease in cell growth compared to the scrambled siRNA-transfected control. Red bars indicate the knockdown of kinases that significantly decreased cell viability. ANOVA, ** indicate p<0.001, * indicate p<0.05. **E.** Time course cell growth curves of ZFTA-RELA^fus^ ZR1 cells transfected with scrambled siRNA control or siRNAs targeting various kinases: Acvr1 and Acvr1b (top row left), Ptk6 (top row middle), Fgfr1 and Mertk (top row right), Epha1 and Epha6 (bottom row left), Irak1 (bottom row middle), and Pim3 (bottom row right). Cell confluence was assessed as the percentage of the total image area occupied by cells.

These responses were used as input for an elastic net-regularized multiple linear regression model, with the residual activities of 298 kinases serving as explanatory variables. Model performance was evaluated using leave-one-out cross-validation (LOOCV), comparing predicted versus observed growth responses. The optimized model achieved over 90% predictive accuracy. From the best-performing models, we identified 12 kinases most strongly associated with ZR1 cell growth, including several receptor tyrosine kinases such as MERTK, members of the FGFR family, c-Kit, PTK6, and EPHA1 (**Fig. 2B**), highlighting potential therapeutic targets in ZFTA-RELA-driven ependymoma.

### Validation of kinase dependencies in ZFTA-RELA-driven ependymoma

To validate the accuracy of our KiR predictions, we performed siRNA-mediated knockdown of the 12 kinases identified as top candidates and assessed their impact on the viability of ZR1 cells. Quantitative PCR confirmed at least a 50% reduction in expression for each targeted kinase compared to non-targeting controls (**Fig. S3**). Knockdown of 9 out of the 12 kinases resulted in a statistically significant decrease in ZR1 cell viability, with Mertk knockdown causing more than a two-fold reduction, highlighting it as a particularly strong candidate (**Fig. 2C**).

To further assess functional impact, we used live-cell imaging to monitor growth following knockdown of Mertk, Fgfr1, Epha1, Epha6, Ptk6, Acvr1, Acvr1b, Pim3, and Irak1. Depletion of Mertk, Epha1/6, Ptk6, Acvr1/b, Pim3, and Irak1 significantly impaired cell growth compared to both untransfected and siRNA control conditions (**Fig. 2D**). While FGFR1 and FGFR3 have previously been implicated in ST-EPN, our identification of IRAK1, known to regulate NF-κB signaling, a hallmark of ZFTA-RELA^fus^ tumors, further supports the accuracy of the KiR approach. Notably, Mertk had not been previously associated with ependymoma, pointing to a novel and potentially actionable vulnerability.

We next asked whether any of these KiR-predicted kinases are aberrantly expressed in human ZFTA-RELA^fus^ tumors. Transcriptomic comparisons between ZFTA-RELA^fus^ tumors, low-grade supratentorial tumors, and YAP-fusion-driven ependymomas revealed significant upregulation of MERTK, ACVR1B, PIM3 in the ZFTA-RELA^fus^ group (**Fig. 3A**). FGFR1 and EPHB2 were upregulated in both ZFTA-RELA^fus^ and YAP-fus tumors. Consistently, analysis of RNA-seq data from 370 ependymoma samples revealed significantly higher expression of MERTK, FGFR1, PIM3, and EPHB4 in the EPN-E1 subtype compared to EPN-E2, reinforcing their selective enrichment in ZFTA-RELA^fus^-driven tumors (**Fig. 3B**).

**Figure 3.**
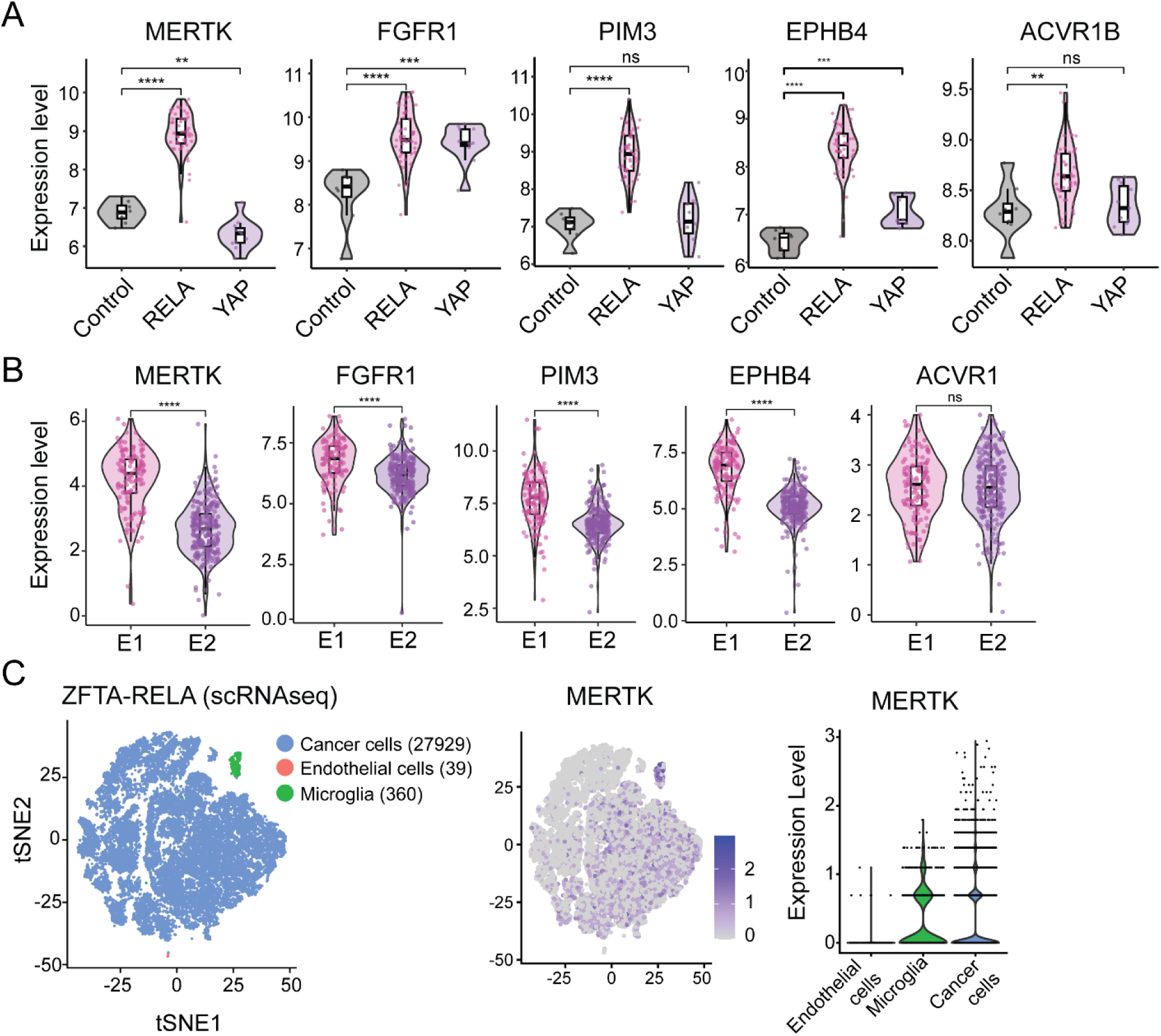
Elevated expression of MERTK and other kinases in ZFTA-RELA^fus^ ependymomas across bulk and single-cell datasets. **A.** Violin plots showing the relative expression of MERTK, FGFR1, PIM3, EPHB4, and ACVR1B in human ependymoma tumors, comparing ZFTA-RELA^fus^, YAP^fus^, and low-grade supratentorial tumors. MERTK, FGFR1, and EPHB4 are significantly upregulated in ZFTA-RELA^fus^ tumors. Significance was determined by Wilcoxon rank-sum test; ****p<0.0001, **p<0.01, ns = not significant. **B.** Expression of the same kinases in EPN-E1 versus EPN-E2 molecular subgroups shows selective enrichment in the EPN-E1 cluster, which is dominated by ZFTA-RELA^fus^ tumors. Significance was determined by Wilcoxon rank-sum test; ****p<0.0001, ns = not significant. **C.** Single-cell RNA-seq analysis of a ZFTA-RELA^fus^ tumor. Left: UMAP plot identifying major cell populations. Middle: MERTK expression mapped across all cells, indicating expression in a subset of tumor cells. Right: Violin plot confirming MERTK expression in ZFTA-RELA^fus^ cancer cells, supporting a tumor-intrinsic role for MERTK in this subtype.

MERTK is well known to be expressed in myeloid cells within the tumor microenvironment, where it promotes M2-like polarization and immunosuppression(*20*). However, its specific role in tumor cells remains less well-defined. While bulk RNA-seq data suggested elevated MERTK expression in ZFTA-RELA^fus^ ependymomas, this does not distinguish between tumor and non-tumor cell types. To clarify this, we analyzed a previously published single-cell RNA-seq dataset from a two ZFTA-RELA^fus^ supratentorial ependymoma patients (*3*), comprising >28,000 cells (**Fig. 3C**). Our analysis confirmed that MERTK is highly expressed in microglia, as expected, but also revealed clear expression in tumor cells, supporting a potential cell-intrinsic role for MERTK in ZFTA-RELA^fus^-driven ependymoma **(Fig. 3C)**. Supporting these transcriptomic findings, we also observed elevated Mertk protein levels in ZFTA-RELA^fus^ mouse tumors compared to naïve brain tissue (**Fig. 4A**). Together, these data validate the functional significance of several KiR-predicted kinases and highlight MERTK as a novel, biologically relevant target in ZFTA-RELA-driven ependymoma.

**Fig. 4.**
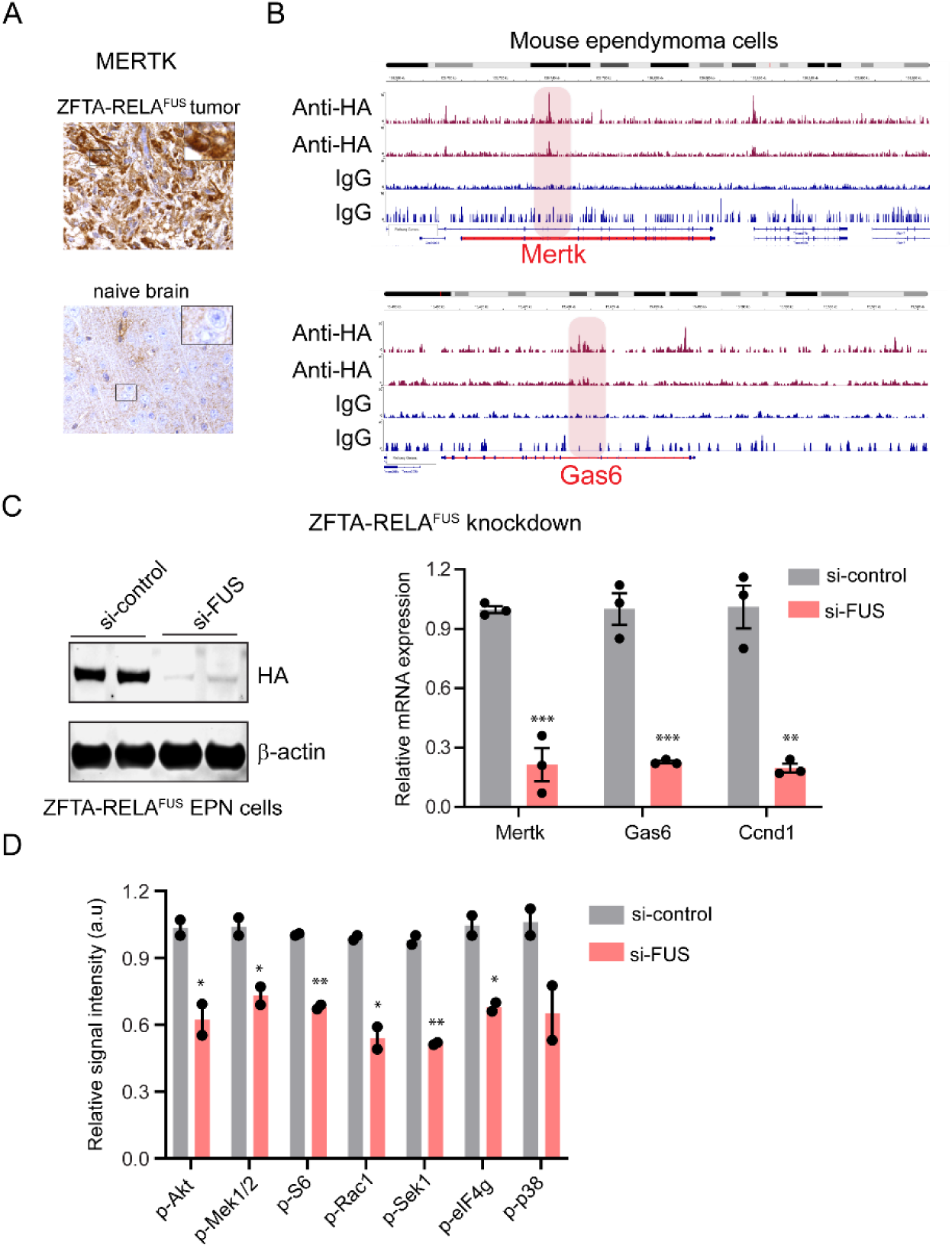
ZFTA-RELA^fus^ drives the expression of Mertk and Gas6. **A.** Representative images showing protein expression of Mertk in ZFTA-RELA^fus^ mouse tumor and naïve brain detected via immunostaining. **B**. ZFTA-RELA^fus^ binding sites were identified in the promoter regions of Mertk and Gas6 in mouse EPN cells. Anti-HA was used to pull down the ZFTA-RELA^fus^, while IgG was used as a negative control. **C.** Knockdown of ZFTA-RELA^fus^ caused downregulation of Mertk, Gas6, and cyclin D1 (Ccnd1). *Left*, a Western blot showing the effect of siRNAs targeting RELA on the protein expression of ZFTA-RELA^fus^, as measured by immunoblotting with HA antibody. On the right, the effect of siRNAs targeting ZFTA-RELA^fus^ on Mertk, Gas6, and Ccnd1 was assessed using quantitative PCR, with data normalized to GAPDH expression. Bars represent the mean of three replicates samples, and error bars indicate the SEM. Unpaired, two-tailed, t-test, *** indicate p<0.0001, ** indicate p<0.01. **D**. siRNAs targeting ZFTA-RELA^fus^ inhibits the phosphorylation of several pro-growth and survival signaling proteins. Bars represent the mean of two replicates and error bares represent SEM. Unpaired, two-tailed, t-test, ** indicate p<0.01, * indicate p<0.05.

### ZFTA-RELA^fus^ directly activates autocrine MERTK signaling to promote oncogenic pathways

Previously, it was shown that in addition to activating the NFκB pathway, ZFTA-RELA^fus^ drives a neoplastic transcriptional program by binding to thousands of unique sites across the genome, activating pathways enriched in MAPK and focal adhesion networks (*21, 22*). We investigated whether ZFTA-RELA^fus^could directly activate MERTK signaling. ChIP-seq analysis from two recent EPN studies revealed prominent binding sites for ZFTA-RELA^fus^ in the promoter regions of both Mertk and its cognate ligand, Gas6, in mouse EPN cells (**Fig. 4B**). Further, we observed elevated levels of Gas6 in E1 EPN subcluster compared to E2 subcluster (**Fig. 3B**), suggesting that ZFTA-RELA^fus^ could prime the activation of autocrine Gas6-Mertk signaling in cancer cells. To validate this, we knocked down ZFTA-RELA^fus^ in ZR1 cells and assessed changes in *Mertk* expression (**Fig. 4C**). Our data showed that the knockdown of ZFTA-RELA^fus^ led to a significant decrease in the mRNA expression of *Mertk*, *Gas6*, and *Ccnd1* (cyclin D1, a known ZFTA-RELA^fus^ downstream target gene) (**Fig. 4C)**. Furthermore, knockdown of ZFTA-RELA^fus^ resulted in decreased phosphorylation of several pro-growth and survival signaling proteins, including Akt, S6 ribosomal protein, eIF4G, and Mek-Erk (**Fig. 4D)**. These findings suggest that ZFTA-RELA^fus^ not only activates the NFκB pathway but also promotes Mertk signaling by directly binding to its promoter region, highlighting a dual mechanism by which ZFTA-RELA^fus^ drives ependymoma progression.

Previous studies have shown that Mertk regulates intracellular pro-survival and anti-apoptotic signaling pathways in cancer cells and skews macrophage polarization towards a pro-tumor M2-like phenotype in the tumor microenvironment (*20, 23, 24*). To investigate the contribution of Mertk activation in ZFTA-RELA^fus^-expressing cancer cells, we stimulated ZR1 cells with Gas6 and observed a robust increase of the phosphorylation of Erk1/2, Akt and S6 ribosomal protein within 5 minutes, indicating that Gas6-Mertk signaling is functional and intact in ZFTA-RELA^fus^ cancer cells (**Fig. 5A**). Given that Gas6 can also activate other members of the TAM receptor family, including Axl and Tyro3, we sought to confirm that the activation of the MAPK and AKT pathways in response to Gas6 is specifically due to Mertk. We treated ZR1 cells with the Mertk inhibitor MRX2843 prior to Gas6 stimulation (**Fig. 5B**). Our data showed that Mertk inhibition significantly blocked the activation of several downstream pathways, including Mek/Erk and Akt/S6 **(Fig. 5B)**, suggesting that the Mertk receptor is a key driver of Gas6-mediated signaling in ZFTA-RELA^fus^ cells.

**Fig. 5.**
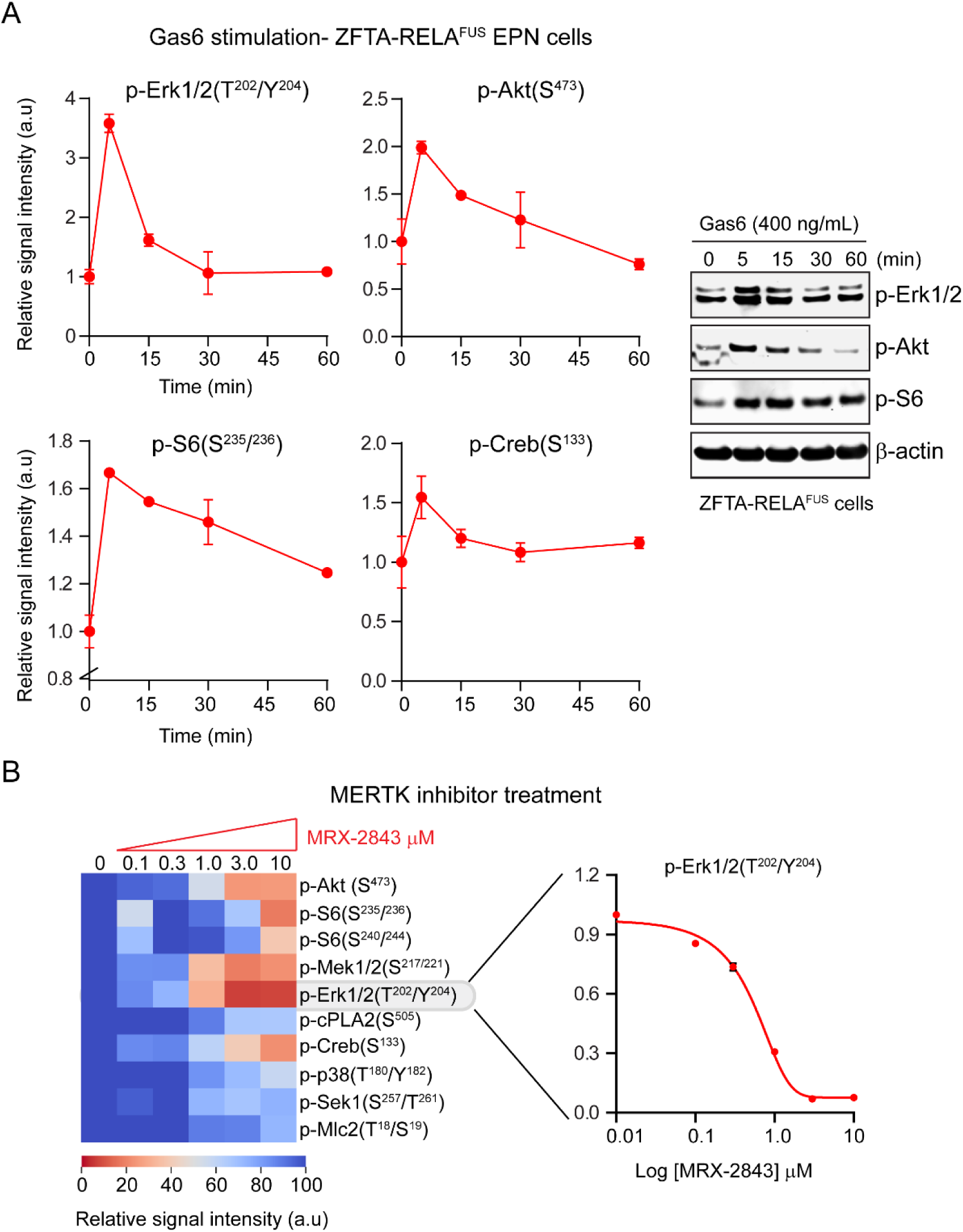
Gas6-Mertk signaling drives pro-growth signaling cascade in ZFTA-RELA^fus^ EPN cells. **A.** Gas6 stimulation caused activation of several growth factor signaling proteins in a time-dependent manner. Plots showing changes in the phosphorylation of Erk1/2, Akt (pan), S6, and Creb in ZFTA-RELA^fus^ EPN cells treated with Gas6. *Right*, representative Western blot images Phosphorylation changes were measured by Western blot. **B**. MRX2843 treatment (1 hour) inhibits several growth factors signaling proteins in a time dependent manner. A heatmap showing changes in the phosphorylation levels of Akt (pan), S6 ribosomal protein, Mek1/2, Erk1/2, cPLA2, Creb, p38, Sek1, and Mlc2 in ZFTA-RELA^fus^ EPN cells treated with the Mertk inhibitor, MRX-2843, at various concentrations (from 0.1 to 10 µM).

### MERTK inhibition suppresses tumor growth and oncogenic pathways in ZFTA-RELA^fus^ EPN

Having established that MERTK signaling is active and contributes to pro-survival pathways in ZFTA-RELA^fus^ ependymoma cells, we next evaluated the therapeutic potential of MRX-2843, a clinical-grade MERTK inhibitor capable of crossing the blood-brain barrier (*25, 26*). In 2D cultures of ZR1 cells, MRX-2843 significantly reduced cell viability, with an EC₅₀ of ∼200 nM (**Fig. 6A**). This effect was consistent across four patient-derived ZFTA-RELA^fus^ cell lines, while minimal sensitivity was observed in ZFTA-MAML^fus^ cells, indicating subtype-specific vulnerability (**Fig. 6B**). MRX-2843 also reduced viability of ZFTA-RELA^fus^ tumor slices *ex vivo* in a dose-dependent manner (**Fig. 6C**).

**Fig. 6.**
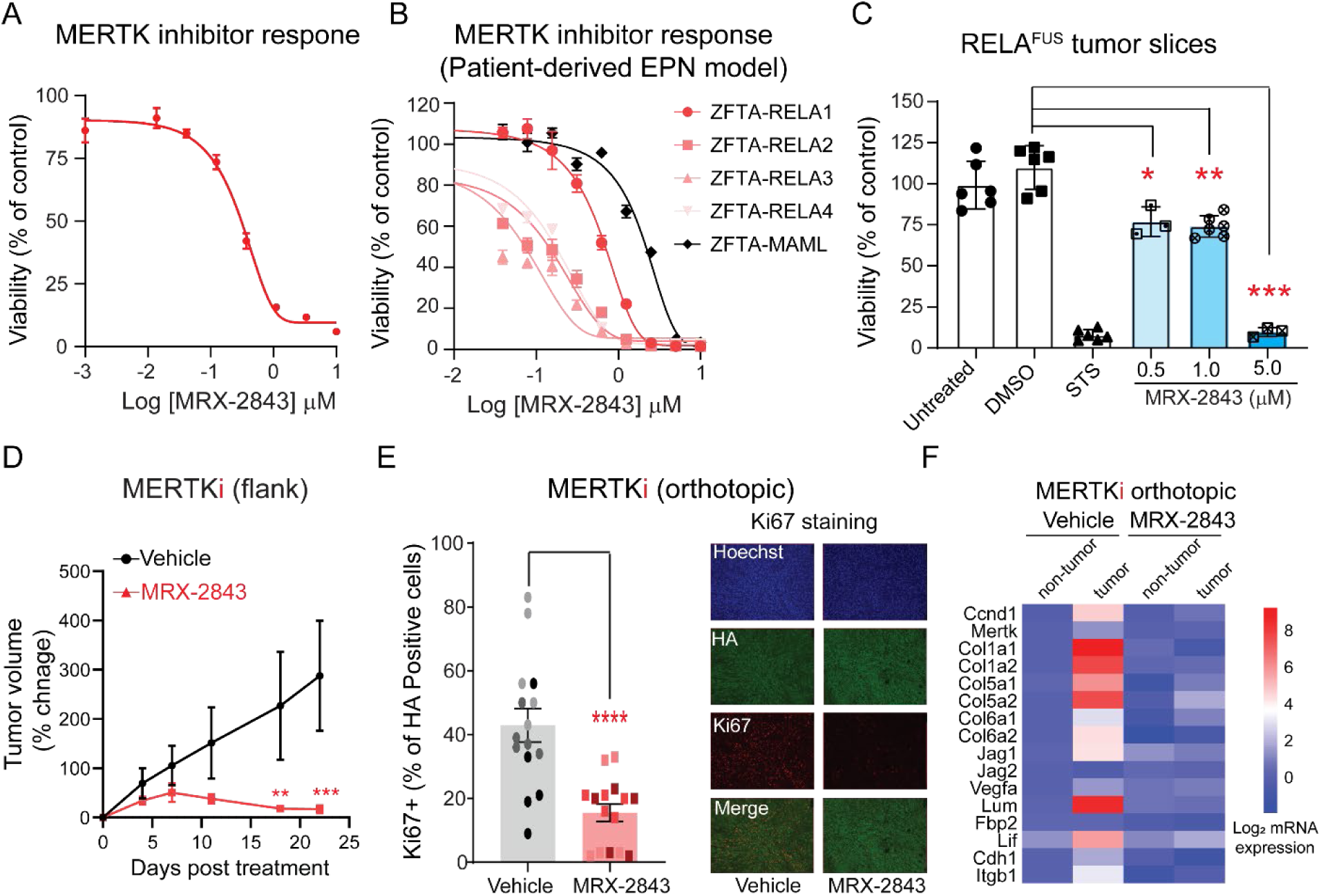
MERTK inhibition suppresses tumor growth in ZFTA-RELA^fus^ ependymoma. **A.** ZFTA-RELA^fus^ mouse tumor-derived cells (ZR1) exhibit strong sensitivity to the MERTK inhibitor MRX-2843 in 2D culture, with an EC₅₀ of ∼200 nM. **B.** MRX-2843 selectively inhibits viability in four patient-derived ZFTA-RELA^fus^ cell lines, but not in a ZFTA-MAML^fus^ control line, indicating subtype-specific sensitivity. **C.** *Ex vivo* treatment of ZFTA-RELA^fus^ tumor slices with MRX-2843 results in a dose-dependent reduction in tissue viability. Staurosporine (STS) was used as a positive control. *p<0.05, **p<0.01, ***p<0.001 by Student’s t-test. **D.** MRX-2843 significantly reduces tumor growth in mice bearing subcutaneous ZR1 flank tumors over a 21-day treatment period. **E.** Short-term (5-day) MRX-2843 treatment in mice with orthotopic ZR1 tumors reduces proliferation, as indicated by decreased Ki67 staining in HA-positive tumor cells. Right, representative immunofluorescence images showing Ki67, HA, and Hoechst staining on ZFTA-RELA^fus^ tumors, captured at 20x magnification with a scale bar of 50 µm. **E.** Heatmap showing changes in the expression of ZFTA-RELA^fus^ EPN marker genes in response to short-term MRX-2843 treatment in mice. Data are normalized vehicle-treated non-tumor tissue.

*In vivo*, MRX-2843 treatment significantly reduced tumor growth in mice bearing subcutaneous ZR1 flank tumors (**Fig. 6D**). Mice were treated once daily for 21 days, and MRX-2843 led to a clear reduction in tumor volume compared to vehicle-treated controls, with no significant weight loss or signs of toxicity, indicating good tolerability (**Fig. 6D**). Given that the orthotopic mouse model of ZFTA-RELA^fus^ ependymoma exhibits high variability in tumor growth, making it challenging to perform long-term drug efficacy studies, we focused on evaluating early pharmacodynamic effects. Mice harboring intracranial ZR1 tumors were treated with MRX-2843 for 5 days, followed by assessment of tumor proliferation via Ki67 staining. This short-term treatment led to a marked reduction in proliferative HA+ tumor cells compared to vehicle-treated controls (**Fig. 6E**), suggesting that MERTK inhibition suppresses tumor cell proliferation in the native brain microenvironment. Molecular analysis of MRX-2843-treated ZFTA-RELA^fus^ tumors compared to vehicle-treated controls revealed a substantial downregulation of key fusion marker genes and tumor-associated pathways. Specifically, treatment with MRX-2843 led to the complete suppression of the Ccnd1, which is critical for cell cycle progression in EPN (*27*). In addition, genes involved in extracellular matrix (ECM) remodeling, such as various collagens, Lumican (Lum), and Integrin β1 (Itgb1), were significantly downregulated, indicating a disruption in tumor structural integrity and invasive potential (**Fig. 6E**). Furthermore, inflammatory signaling genes, particularly Lif, known to contribute to tumor progression and immune evasion(*28*), were also markedly reduced following MRX-2843 treatment. Together, these data suggests that Mertk plays an important role in the growth of ZFTA-RELA^fus^ cells and tumors by disrupting multiple essential regulators of ependymoma progression, including extracellular matrix remodeling and immune evasion genes.

To gain a comprehensive understanding of the pathways and processes affected by Mertk inhibition in ZFTA-RELA^fus^ EPN tumors, we conducted unbiased transcriptomic profiling. Short-term treatment (5 days) with MRX-2843 resulted in significant gene expression changes, with 3,868 genes downregulated and 3,514 genes upregulated compared to vehicle-treated tumors (**Fig. 7A**). Notably, there was substantial overlap between genes upregulated in ZFTA-RELA^fus^ tumors and those downregulated following MRX-2843 treatment, indicating that Mertk inhibition directly disrupts tumor-promoting pathways (**Fig. 7B**). Mertk inhibition also decreased *Zfta* expression, a downstream target of the ZFTA-RELA^fus^, suggesting a feed-forward loop that promotes tumor progression (**Fig. 7C**). Gene Set Enrichment Analysis (GSEA) of significantly downregulated genes following Mertk inhibitor treatment revealed strong enrichment of pathways related to epithelial-mesenchymal transition (EMT), interferon signaling, and inflammatory responses (**Fig. S2**). Consistently, Gene Ontology analysis of the downregulated genes revealed significant enrichment in pathways related to ECM organization, and cytokine signaling, while the upregulated genes were enriched in pathways associated with neuronal systems and GPCR signaling (**Fig.7D**). Finally, PROGENy pathway analysis(*29*) comparing naïve brain tissue and ZFTA-RELA^fus^ tumors revealed significant enrichment of pro-oncogenic pathways, including WNT, JAK-STAT, and NF-κB signaling (**Fig.7E**). Notably, all these pathways were markedly downregulated in Mertk inhibitor-treated ZFTA-RELA^fus^ tumors. Collectively, these findings highlight the therapeutic potential of targeting MERTK with MRX-2843 in ZFTA-RELA^fus^-driven ependymomas, demonstrating a substantial impact on tumor growth, viability, and the suppression of critical oncogenic pathways.

**Fig. 7.**
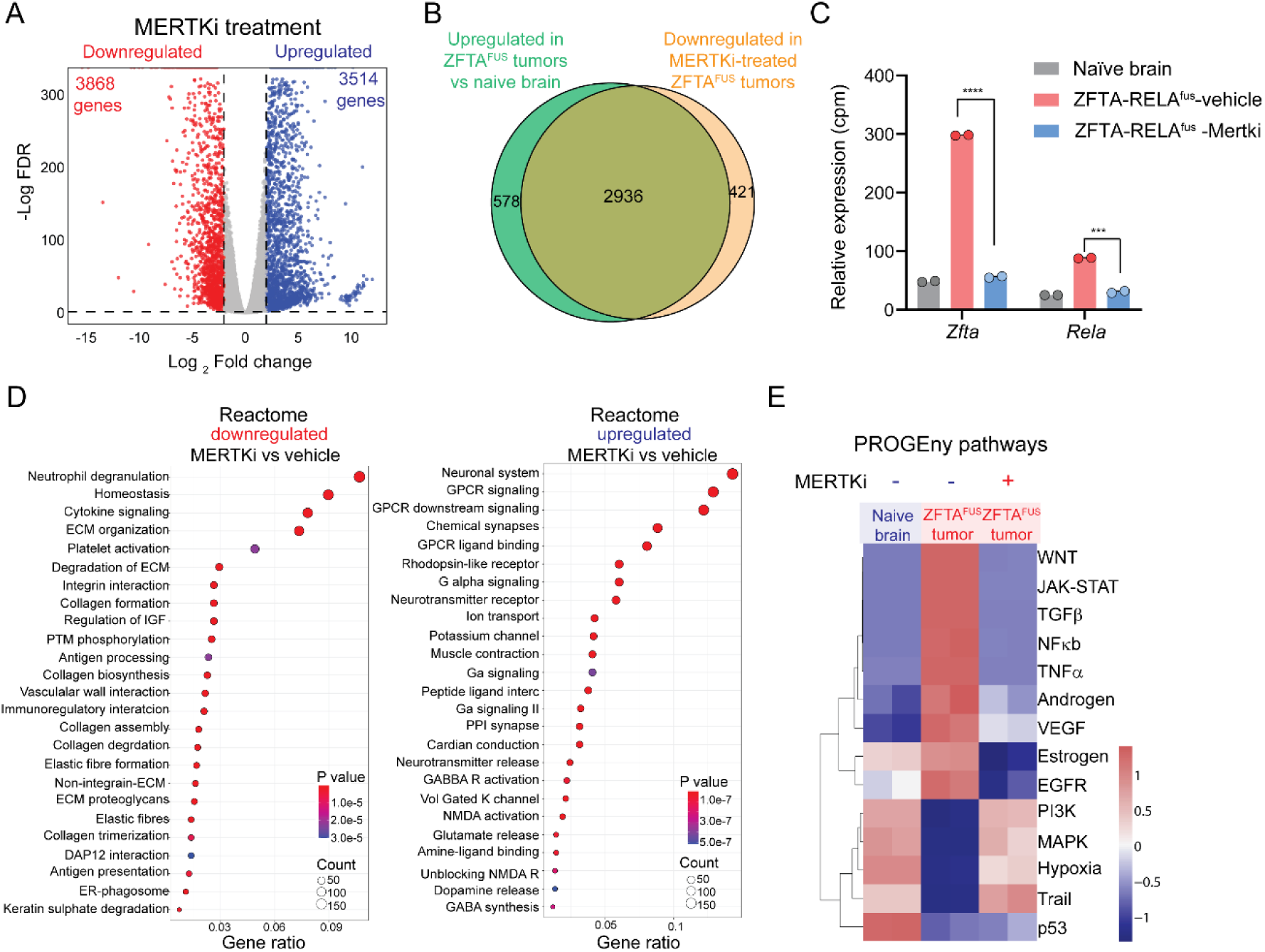
Mertk inhibition suppresses oncogenic programs and alters transcriptional networks in ZFTA-RELA^fus^ ependymomas. **A.** Volcano plot showing differentially expressed genes in ZFTA-RELA^fus^ tumors compared to naïve brain tissue. A total of 3,868 genes were significantly downregulated (red) and 3,514 genes were significantly upregulated (blue). **B.** Venn diagram illustrating the overlap between genes upregulated in ZFTA-RELA^fus^ tumors versus naïve brain tissue (green) and genes downregulated in Mertk inhibitor-treated ZFTA-RELA^fus^ tumors (orange). A total of 2,936 genes were commonly altered, indicating that MERTK inhibition disrupts tumor-promoting pathways. **C.** Mertk inhibition decreased the expression of both *Zfta* and *Rela* in ZFTA-RELA^fus^ tumors. **D.** GO analysis significantly downregulated (left) and upregulated (right) pathways after Mertk inhibition. Pathways are ranked based on significance (*P* value) and gene count, with colors indicating statistical significance. **E.** A heatmap depicting PROGENy pathway analysis showing significant enrichment of pro-oncogenic pathways (WNT, JAK-STAT, and NF-κB) in ZFTA-RELA^fus^ tumors, all of which are markedly downregulated after MERTK inhibition.

## Discussion

Broadly, targeting specific pathways in cancer is challenging due to the presence of multiple concurrent and redundant drivers, especially in primary cancers of the central nervous system, where the blood-brain barrier further limits treatment options. However, some tumors driven by single events essential for tumor survival, such as chromosomal translocations or rearrangements resulting in gene fusions, can be effectively targeted by drugs aimed at these genetic aberrations. For example, the BCR-ABL fusion in chronic myeloid leukemia and the corresponding success of imatinib provide robust and sustained therapeutic targets(*30*). Similarly, the ZFTA-RELA^fus^ is a singular event that drives ependymomas and thus could be targeted by agents that directly inhibit the fusion protein. However, directly targeting the ZFTA-RELA^fus^ protein, a constitutively active transcription factor, is challenging. Therefore, targeting proteins downstream of the RELA fusion oncoprotein that are essential for tumor transformation represents an attractive alternative. Kinase inhibitors, with more than 80 kinases clinically approved, represent one of the major classes of anti-cancer drugs(*31*). Identifying kinases downstream of the ZFTA-RELA^fus^ could accelerate clinical translation and offer a promising strategy for targeted therapy.

In this study, we identified two molecular subtypes of ependymoma, EPN-E1 and EPN-E2, each distinguished by unique gene fusions, kinase expression patterns, and pathway activity. The EPN-E1 cluster, which includes the majority of ZFTA-RELA^fus^ supratentorial tumors, represents a clinically aggressive subtype with limited treatment options. Our comparative transcriptomic analysis revealed that the ZFTA-RELA^fus^ mouse model recapitulates the gene expression signature of the human EPN-E1 subtype with high fidelity (**Fig. 1**). This strong concordance establishes the ZFTA-RELA^fus^ model as a biologically relevant and translationally valuable system for uncovering subtype-specific therapeutic targets and advancing drug development for patients with high-risk EPN-E1 tumors.

We utilized ZFTA-RELA^fus^ mouse model and applied a systems pharmacology-based approach to identify druggable kinases downstream of ZFTA-RELA^fus^ in ST-EPN, offering promising opportunities for developing effective treatments for these aggressive tumors. We identified several kinases, including Mertk, Epha1/6, Fgfr1, Ptk6, and Acvr1, as critical for the growth and survival of ZFTA-RELA^fus^ driven ependymoma cells in culture. Specifically, we observed strong inhibition of ZFTA-RELA^fus^-driven cell growth in response to the depletion of Mertk, Ptk6, Epha1, and Acvr1 (**Fig. 2**). Among these, we extensively validated the role of Mertk in various experimental settings, including cultured mouse and patient-derived cells, tumor tissue slices, and a mouse model (**Fig. 6**). Previous studies have implicated Fgfr1 in the growth of aggressive ependymomas(*32, 33*); however, the roles of the other kinases remain to be fully investigated. Notably, the inhibition of Mertk in tumors led to a significant decrease in the expression of ECM components, such as collagens and integrins (**Fig. 6, 7**). Both Acvr1, a member of the TGF-beta family, and Eph receptors are known to regulate ECM components in the tumor microenvironment(*34–36*). Furthermore, recurrent ACVR1 mutations have been observed in posterior fossa ependymomas(*37*). It is conceivable that the kinases identified in our screening approach could work together to regulate key biological processes in ependymoma progression. Therefore, an approach that involves targeting a combination of these kinases could be explored for more effective treatments.

Mertk, a member of the TAM family of receptor tyrosine kinases, is aberrantly expressed in several cancers, including central nervous system tumors such as astrocytoma(*38*) and glioblastoma(*39*). Mertk signaling contributes to an immunosuppressive tumor microenvironment by inducing an anti-inflammatory cytokine profile, and modulating the functions of macrophages, myeloid-derived suppressor cells, natural killer cells, and T cells(*40*). Consistently, our data show that short-term inhibition of Mertk in tumors led to a significant decrease in the immunosuppressive cytokine Lif. Notably, Mertk inhibition also reduced *Zfta* expression, which is also a downstream target of the ZFTA-RELA^fus^, suggesting the presence of a feed-forward loop that amplifies tumor progression (**Fig. 7C**). This feed forward loop cannot occur in the mouse model as the ZFTA-RELA fusion gene is driven by the RCAS LTR promoter rather than the ZFTA promoter as in the human case. Disrupting this loop through Mertk inhibition could completely abrogate tumor growth in human ZFTA-RELA^fus^ ependymomas. Furthermore, the substantial overlap (>80%) between genes upregulated in ZFTA-RELA^fus^ tumors and those downregulated after MRX-2843 treatment (**Fig. 7B**) supports that Mertk inhibition directly disrupts tumor-promoting pathways. Therefore, targeting Mertk in ZFTA-RELA^fus^ EPN could serve a dual purpose: directly eliminating cancer cells and transforming the microenvironment to support anti-tumor activity. Collectively, these findings highlight the therapeutic potential of targeting Mertk with MRX-2843 to suppress tumor growth and block key oncogenic pathways in ZFTA-RELA^fus^-driven ependymomas.

In conclusion, our findings demonstrate that the ZFTA-RELA^fus^ mouse model effectively replicates human ependymoma subtype. By combining this model with a systems pharmacology-based approach, we identified Mertk as a critical kinase involved in the progression of ZFTA-RELA^fus^-driven ependymomas. The significant impact of Mertk inhibition on tumor proliferation in preclinical models suggests that targeting this kinase could be an effective therapeutic strategy for patients with RELA fusion-driven ependymomas. These findings pave the way for further clinical evaluation of Mertk inhibitors, offering hope for improved treatments for this challenging and aggressive cancer type.

## Materials and methods

### Human bulk RNASeq analysis Collection of publicly available RNA Sequencing data

Raw RNA sequencing data for ependymoma samples were retrieved from various public data repositories, as detailed in Table S1. The RNA sequencing reads were then aligned to the Gencode GRCh38.primary_assembly reference genome using STAR(*41*) (v2.7.7a). Gene-level quantification was conducted with HTSeq(*42*) (v0.11.0) using Gencode(*43*) V39 primary assembly annotations. The raw gene counts from all datasets were subsequently aggregated and batch effects were corrected using the ComBat-seq function from the R package “sva”(*44*). Normalized gene expression values were calculated and expressed as VST from “DESeq2”(*45*) package. consensusClusteringPlus(*46*) was used to cluster the samples.

Gene fusions were identified using the Arriba(*47*) tool (v2.1.0) and STAR-Fusion(*48*) on RNA-Seq reads aligned via STAR’s two-pass method. Fusion analysis was restricted to high-confidence fusions found by both Arriba and STAR-Fusion sing only protein-coding genes. All figures were made in R, using ggplot2(*49*).

### Mouse modeling of ZFTA-RELA^fus^ using RCAS/tva

The RCAS/TV-A system was employed for mouse studies(*50*). RCAS vector ZFTA-RELA^fus^-HA was transfected into DF-1 cells. Neonatal mice of the Ntva, Ink4a/Arf, aged between 3 and 5 days, received an intercranial injection of 100,000 cells. Tumor formation was confirmed by weight loss, poor grooming leading to MRI imaging at approximately 2-3 months of age.

### Mouse ZFTA-RELA^fus^ cells and reagents

The mouse ependymoma cell line ZR1 was established from a ZFTA-RELA^fu^**^s^** tumor by dissociating the tumors and culturing the cells short-term. The presence of the ZFTA-RELA^fus^ fusion in the ZR1 cells was confirmed through immunofluorescence staining using an HA antibody. The cells were cultured in DMEM supplemented with 10% (v/v) fetal bovine serum (FBS), 100 IU/mL penicillin, and 100 µg/mL streptomycin. Kinase inhibitors were purchased from Selleck Chemicals (Houston, TX, USA), or Cayman Chemical Inc. (Ann Arbor, MI, USA). MRX-2843 for animal studies was obtained from MedChemExpress (Monmouth Junction, NJ, USA). Kollisolv PEG E 400, and pharmaceutical grade DMSO were purchased from Sigma-Aldrich. Normal saline was obtained from Fisher Scientific (Waltham, MA, USA). Sodium sulphobutylether-beta-cyclodextrin was acquired from AK Scientific (Union City, CA, USA).

### Preparation of tissue slices

Organotypic tumor slices were prepared from mouse ependymoma tumors as described previously (*51*). Fresh tumor tissues were shaped into 6 mm diameter cylinder using biopsy punches (Integra Miltex, York, PA, USA). The cores of the tumor were sliced at 250 μm thickness using a Leica VT1200 S vibrating blade microtome (Leica Biosystems, Wetzlar, Germany). The tissue cores were mounted using cyanoacrylate glue and submerged in ice-cold HBSS during slicing. The slicing parameters included a blade angle of 18°, a speed of 0.15 mm/s (with a range of 0.10 to 0.28) mm/s, and vibration amplitude of 2.0 mm. Slices were cultured on air-liquid interface using cell culture inserts in slice culture medium containing Williams’ Medium supplemented with 12 mM nicotinamide, 150 nM ascorbic acid, 2.25 mg/mL sodium bicarbonate, 20 mM HEPES, 50 mg/mL glucose, 1 mM sodium pyruvate, 2 mM L-glutamine, 1% (v/v) ITS+Premix, 20 ng/mL EGF, 40 IU/mL penicillin, and 40 μg/mL streptomycin.

### Viability measurements of tissue slices and drug screening

Tissue slices were incubated with Realtime-Glo MT Cell Viability reagents (Promega, Madison, WI, USA), with 50 μL medium on top. Two days from preparation, drugs were added to the tissue slices at final concentration at 0.5 µM, 1.0 µM or 5.0 µM. The slices were incubated for at least 2 hours to overnight with agitation at 100-150 rpm on an orbital shaker in a cell incubator, followed by a static incubation. Bioluminescence from tissue slices was measured using a BioTek Syngery H4 Hybrid microplate reader (BioTek, Winooski, VT, USA). Viability was calculated by the ratio of the signal at day 5 to the signals from the same tissue before treatment, then normalized to the value of vehicle control slices to be 100%.

### Kinetic in vitro cell growth assay

ZR1 cells or its variants with siRNA transfection were seeded at 5,000 cells in 96-well plates. The confluence of the cells to the imaged area was evaluated using IncuCyte Live-Cell Imaging System and accompanying cell analyzer software (Essen BioScience, Ann Arbor, Michigan, USA). The inhibitor effect was normalized to the control when the control wells reached 70% confluence. To evaluate the effect of Mertk inhibitor, MRX-2843, 5,000 cells were seeded in 96-well plates, and cell viability was measured at 72-96 hours using CellTiter-Glo 2.0 reagents (Promega, Madison, WI) following the manufacturer’s instructions.

### Small interfering RNA transfection

All small interfering RNAs (siRNAs) were obtained from Dharmacon (Lafayette, CO, USA). siRNA’s transfections in 6-well plates for expression profiling and in 96-well plates for monitoring changes in cell growth were carried out using Lipofectamine RNAiMax (Invitrogen, Waltham, MA, USA) according to the manufacturer instructions.

### RNA extraction and quantitative PCR

Total cellular RNA was isolated using a GenCatch Total RNA Mini Prep Kit (Epoch Life Science, Missouri City, TX, USA). mRNA expression changes were determined using quantitative real-time PCR (qPCR). Briefly, 1 μg of total RNA was reverse transcribed into first-strand cDNA using an RT2 First Strand Kit (Qiagen, Hilden, Germany). The resultant cDNA was subjected to qPCR using gene-specific primers (BioRad, Hercules, CA, USA) and GAPDH (housekeeping control). The qPCR reaction was performed using a BioRad CFX384 thermocycler with an initial denaturation step of 10 min at 95 °C, followed by 40 cycles of 15 s at 95 °C and 60 s at 58 °C. The mRNA levels of genes encoding cytokine expression were normalized relative to the mean levels of the housekeeping gene and compared using the 2^−ΔΔCt^ method as described previously (*52*).

### Western blotting

Monolayer cells were washed twice with PBS and suspended in 2% SDS lysis buffer. The remaining debris were removed by centrifuge filtration. Protein concentration was measured using a Pierce BCA protein assay kit (Thermo Scientific, #23225) with bovine serum albumin as a standard. After denaturation at 95℃ for 5 min in reduced condition, 10-20 µg of proteins per lane were separated with 4-12% Bis-Tris gradient gels in MES SDS running buffer using the XCell SureLock Mini-Cell Electrophoresis System. The proteins were transferred to nitrocellulose membrane by a BioRad wet/tank blotting system. The membrane was then blocked with Licor Intercept (PBS) blocking buffer (LICOR, #927-70001) for 1 hour at ambient temperature, followed by primary antibody at 1:1000 in the blocking buffer at 4℃ for overnight. Primary antibodies used were as follow: β-actin (Sigma, A1978), HA-tag (Cell Signaling Technology, CST#3724), phospho-Akt (Cell Signaling Technology, CST#4058), phospho-p44/42 MAPK (Erk 1/2) (Cell Signaling Technology, CST#4376), phospho-S6 Ribosomal protein (Cell Signaling Technology, CST#2211), and phospho-CREB (Cell Signaling Technology, CST#9198). Following PBS washes, the membrane was incubated with IRDye 800CW goat anti-Mouse IgG secondary antibody (LICOR # 926-32210) or IRDye 680RD goat anti-Rabbit IgG secondary antibody (LICOR #926-68071) at 1:5000 for 1 h at room temperature. Signal detection was performed using the Odyssey imaging system, and the band intensities were quantified with the Odyssey imaging analysis software. Signal intensities of each protein band were normalized to that of β-actin.

### Reverse-phase protein array

Molecular measurements using Reverse-phase protein arrays (RPPA) were carried out as described previously (*53, 54*). Briefly, mouse ependymoma ZR1 cells treated with various concentration of MRX-2843 were washed twice in PBS, and cell lysates were prepared in 2% SDS lysis buffer as described previously (*54*). Whole cell lysates were printed onto 16-pad nitrocellulose coated slides (Grace Biolabs, GBL505116) using an Aushon 2470 microarrayer (Aushon BioSystems), with each sample printed in triplicate and slides were stored at −20°C until processing. For in vivo samples, ependymoma tumors were surgically extracted from mice brain and cryopreserved until use. Small pieces of the tumors were homogenized and lysed in 2% SDS lysis buffer using a bead ruptor 12 homogenizer (Omni International 19-050A). These samples were processed similarly as to the cell lysate. The tumor lysates were adjusted to 2 mg/mL protein concentration and printed onto nitrocellulose pads attached to slide glass, with 2 depositions with single spot/sample. RPPA slides were washed with 1 M Tris-HCl (pH 9.0) for 2-4 days to remove excessive SDS residue, followed by three washes with PBS for 10 min each. Subsequently, the slides were blocked with Licor Intercept (PBS) blocking buffer for one hour at room temperature. After blocking, arrays were incubated with primary antibodies overnight at 4°C. The next day, arrays were washed thrice with PBS and then incubated with IRDye labeled secondary antibodies in the blocking buffer for 1 hour at room temperature. Following incubation, slides were scanned using a Licor Odyssey CLX Scanner (LICOR). Total signal intensity from each spot was quantified using Array-Pro analyzer software package (Media Cybernetics). The measurement of a specific protein from an individual sample was normalized to its total β-actin (Sigma, A1978).

### Drug treatment of EPN bearing mice

Mice with confirmed tumors by MRI were randomly assigned to receive either a vehicle or 50 mg/kg of MRX-2843. Treatment was administered once daily via oral gavage for either 3 or 5 consecutive days. Mice were euthanized one hour after the final treatment, and the whole brains were harvested for further analysis.

### RNASeq sequencing and analysis

Total RNA was extracted from naïve brain tissue and ZFTA-RELA^fus^ rumors using the RNeasy Mini Kit (QIAGEN). RNA was quantified and assessed for quality using a Nanodrop spectrophotometer (Thermo Fisher). Library prep and RNA sequencing was performed and analyzed at Novogene using a standing workflow with sample quality control (assessed through Library QC) and polyA enrichment. Sequencing was then performed using 150 bp paired-end reads on the Illumina PE150 platform. Raw counts for samples were obtained by aligning sequencing reads to the mm10_assembly reference genome using the STAR aligner (v2.7.7a)(*41*). The aligned files were then indexed with SAMtools (v1.19.2), and gene expression levels were quantified using featureCounts. All raw counts were normalized using log transformed TPM (Transcripts Per Million) values. Differential gene expression analysis was performed between ZFTA-RELA^fus^ vs Naïve brain mouse tumors, using the R/Bioconductor package DESeq2(*45*) (v 1.44.0and a statistical cutoff of FDR < 0.05 and FC > 1.5 was applied to obtain differentially regulated genes. Raw data files have been deposited to the NCBI Gene Expression Omnibus under accession number GEO: GSE000000. GO and Reactome Pathway enrichment analysis was done using R Bioconductor Packages enrichR(*55*) (v 3.2) and dot plots were made using R Bioconductor package ggplot2. Heatmaps were made using R package pheatmap(*56*) (v 1.0.12). Previously published RNAseq analysis. Analysis of RNA-seq microarray expression data were downloaded from the Gene Expression Omnibus (GSE64415). CEL files were processed using the affy package(*57*) in R, with raw data read via the ReadAffy() function and normalized using the Robust Multi-array Average (RMA) method. Gene expression values were mapped using the hgu133plus2 chip annotation. scRNA-seq 10X expression data was obtained from GSE141460. The data was analyzed using the Seurat R package(*58*) (v4.4.0). For each patient sample, cells were filtered to include only those with a total feature count greater than half and less than twice the mean feature count, and with less than 5% mitochondrial (MT) gene content. Normalization was performed using the NormalizeData() function with a scaling factor of 10,000, followed by identification of the top 2,000 highly variable genes using the ‘vst’ method. The data was then scaled and subjected to principal component analysis (PCA) for dimensionality reduction. Clustering was conducted using the first 19 principal components, employing the Louvain algorithm on a shared nearest neighbor (SNN) graph with a resolution parameter of 0.5. Patient samples were subsequently integrated using Harmony(*59*) to produce a unified dataset. Non-linear dimensionality reduction was performed using t-distributed stochastic neighbor embedding (t-SNE) for visualization. Marker gene expression was visualized using Seurat functions (FeaturePlot() and VlnPlot()). All plots were generated using the ggplot2 package (v3.4.4).

### Immunohistochemistry

Harvested brain samples were fixed in 10% buffered formalin phosphate for 48 hours, followed by coronal slicing and processing into paraffin-mounted blocks. Sections of 5 μm thickness were cut and mounted onto charged slides for staining. Hematoxylin and eosin (H&E) staining were performed for tumor verification. Subsequently, DAB antibody staining using a Ventana system was conducted. Stains targeting HA and Ki67 were applied to evaluate proliferation effects on tumor cells with MRX-2843 treatment.

### Immunofluorescence

Immunofluorescent (IF) staining was performed to co-localize HA and Ki67 using Alexa 488 and Cy3, respectively, while Hoechst stain was used for visualizing cell nuclei. This enabled the assessment of the percentage of ZFTA-RELA^fus^ positive cells also positive for Ki67. QuPath software was utilized for IF stain analysis. Each tumor was divided into five 0.5 mm x 0.6 mm sections at 20x magnification, covering the majority of the tumor. Positive counts for ZFTA-RELA^fus^ positive cells were organized into positive and negative counts based on co-staining for Ki67. Minimum intensity thresholds were applied before overlaying ZFTA-RELA^fus^ positive cells with Ki67 positive cells to ensure accurate identification of all positive cells. Positive counts in each section were then expressed as a percentage of all ZFTA-RELA^fus^ positive cells and compared across treatment groups.

## Notes

### Competing Interest Statement

The authors have declared no competing interest.

### Summary of Updates

New data in Figures 1, 3 and 6

### Funding

- Ivy Foundation

